# Best of Both Worlds? Optimising Graph-Based Antimicrobial Resistance Gene Profiling in Long and Short-Read Metagenomes

**DOI:** 10.64898/2025.12.25.693921

**Authors:** David Burke James Mahoney, Finlay Maguire

**Author notes:** Address correspondence to Finlay Maguire.

## Abstract

Environmental surveillance using metagenomic sequencing offers a powerful way to track emerging and mobile antimicrobial resistance (AMR) genes and inform public health mitigation strategies. Read-based analysis tools can sensitively detect AMR genes in metagenomes but provide little information about the surrounding genome. This prevents easily linking detected genes with particular host species or mobile genetic elements. On the other hand, contig-based analysis tools can provide this genomic context but systematically fail to recover many AMR genes. Querying the intermediate assembly graph directly may provide a trade-off between these strengths and weaknesses. However, many existing tools capable of querying assembly graphs are designed for applications other than gene detection, such as pan-genomics, indexing, or scaffolding. Therefore, a comprehensive evaluation of six tools across four search paradigms was performed to determine the optimal graph querying tool for profiling AMR genes in both long and short-read metagenomic assembly graphs. Across mock and simulated metagenomes of varying complexity and read-type, BLAST-based graph alignment (as implemented by GraphAligner) consistently outperformed other graph alignment algorithms. Overall, graph-based methods correctly identified 21% to 46% more AMR genes in complex datasets than contig analyses; however, increases in recall were modest. Combining assembly graphs analyses with contig-based analyses identifies up to 56% additional AMR genes across both long and short-read datasets. This study highlights the challenges associated with metagenomic AMR surveillance and demonstrates that graph-based analyses offer a useful tool in maximising sensitive identification of AMR genes and their genomic context from these data.

**Importance:** Antimicrobial resistance (AMR) is a severe public health threat that has spurred non-governmental organisations and public health agencies to develop action plans to reduce resistance to critical antimicrobials. Surveillance of One Health environments for AMR determinants are often central parts of these action plans. Metagenomic sequencing presents a key method for clinical and public health AMR surveillance; however, algorithmic and biochemical limitations prevent linking most detected AMR genes to their associated host bacteria or mobile genetic elements. Our findings suggest that querying the assembly graph alongside assembled contigs can identify more AMR genes than contigs alone while still providing epidemiologically informative flanking sequences. Associating AMR genes with their genomic context greatly expands our ability to assess the risk they pose across different environments. These improvements in metagenomic AMR gene identification make AMR surveillance more effective for public health institutions potentially reducing the harm of resistant infections.

## Introduction

In 2019, 1.27 million deaths were directly attributable to antimicrobial resistance (AMR), more than malaria and HIV (1). Establishing and strengthening AMR surveillance on local, regional, and national levels is a key objective of the WHO Global Action Plan on AMR in order to maintain the effectiveness of current antimicrobials (2). Metagenomic sequencing has been shown to be an effective method to surveil changes in AMR genes across various One Health environments (3). This is particularly important in interface environments such as wastewater, agricultural waste/byproducts, and peri-urban environments where monitoring the dynamics and transfer of AMR genes between sectors is vital to creating effective One Health interventions (4). However, using metagenomics for AMR surveillance poses a number of technical challenges and trade-offs (5). For example, read-based AMR gene detection methods will sensitively detect AMR genes that are present in the sample; however, information about neighbouring genes is absent which prevents linking a given AMR gene to its associated host species and/or mobile element (6). Conversely, short-read metagenomic contig-based methods provide this genomic context but systematically fail to assemble the majority of AMR gene loci associated with mobile elements (7). To address these challenges, methods for directly querying metagenomic assembly graphs for AMR genes have been proposed. These methods try to exploit the intermediate nature of these graphs to more sensitively detect AMR genes than contig-based methods while recovering more of the AMR genomic context than read-based methods (8–10).

Long-read sequencing is also increasingly used in metagenomics. The longer individual reads can capture some of the genomic context surrounding an AMR gene without requiring assembly. This has enabled read-based analyses which link some detected AMR genes with associated mobile elements (11). However, the adoption and sensitivity of long-read metagenomics has been slightly limited by the challenge of achieving sufficient sequencing depth for complex samples with these technologies (12). It is also not currently clear whether the metagenomic assembly of long-reads results in a similar pattern of systematic loss of AMR genes as short-read assembly. If this is the case, methods for directly querying long-read assembly graphs for AMR genes have the potential to provide a similar performance compromise between read-and contig-based methods.

Querying long or short-read assembly graphs directly requires modifications to traditional sequence/homology search algorithms such as BLAST, Burrows-Wheeler Transform (BWT), k-mer matching, and Hidden Markov Models (HMMs) to account for a non-linear reference (13–18). The majority of graph-based tools such as Bifrost, Minigraph, GraphAligner, Bandage, and SPAligner were primarily developed for the alignment of long-reads to short-read graphs for assembly improvement, pan-genome graph construction, or visualisation (13–16, 18). It is not clear whether these tools are suited to the specific problem of detecting AMR genes in metagenomic assembly graphs. Therefore, this study aims to evaluate the performance of GraphAligner, Bandage, SPAligner, Minigraph, Bifrost, and Pathracer in detecting AMR genes within long and short-read metagenomic assembly graphs and evaluate the relative sensitivity of metagenomic assembly graphs as a means of AMR gene identification.

## Results

### Bandage ran the fastest and Bifrost required the least memory

The computational requirements of existing graph querying tools were assessed to evaluate their feasibility for metagenomic AMR surveillance. Across the high complexity, low complexity, mobile element-enriched, and ZymoBIOMICS graphs, Bandage was fastest for AMR gene detection (19.5s - 552.1s) while the HMM-based Pathracer was the slowest (0.868s - 37513.4s) (Figure S1). Similarly, maximum resident set size (RSS) i.e., peak memory usage, was lowest for k-mer-based Bifrost (0.23 GB to 14.12 GB) and highest for HMM-based Pathracer (0.012 GB - 72.28 GB) (Figure S2). In both runtime and memory usage BLAST-based tools (Bandage and GraphAligner) and BWT-based SPAligner used slightly more time and memory than k-mer-based methods (Figure S1, Figure S2).

### BLAST-based graph querying tools outperform other tools for identifying AMR genes in metagenomic assembly graphs

Assembly graph querying tools were evaluated in terms of their precision and recall in detecting AMR genes across 16 short and long-read metagenomic graphs from different assemblers and of varying complexity. Across all tools, mean recall ranged from 0.051 to 0.381 and mean precision from 0.626 to 0.777 (Figure 1). Of the graph querying tools, BLAST-based GraphAligner had the highest median F_1_-score (0.579) across all metagenomes exceeding direct analysis of assembled metagenomic contigs using RGI (0.441) (Figure 2).

**Figure 1.**
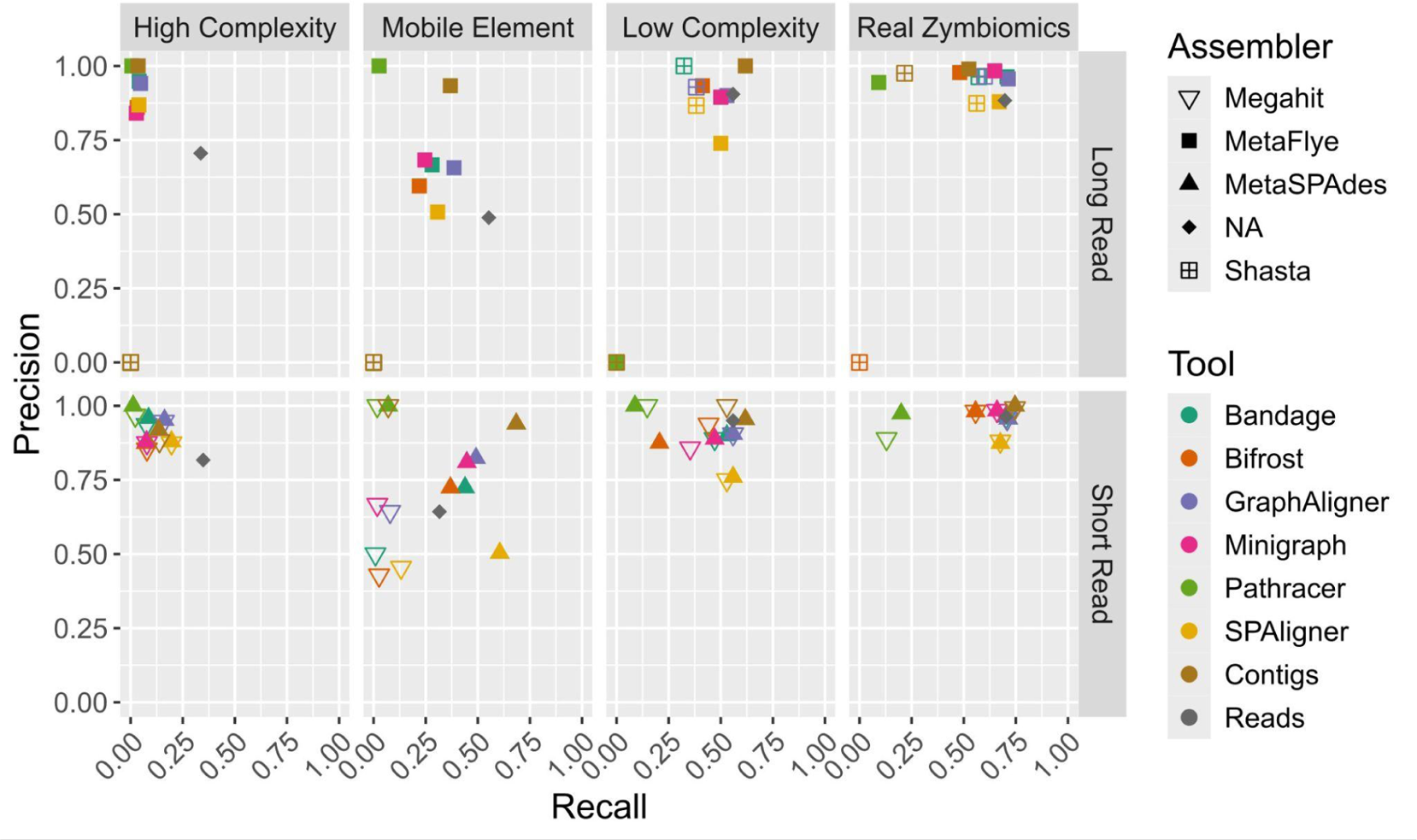
Precision vs. recall of graph querying, contig, and read based tools in identifying the presence of CARD protein homolog model AMR genes for high complexity (6104 taxa), mobile element-enriched (30 taxa), low complexity (29 taxa) and ZymoBIOMICS (10 taxa) long and short-read datasets assembled using MetaSPAdes, MetaFlye, Megahit, and Shasta.

**Figure 2.**
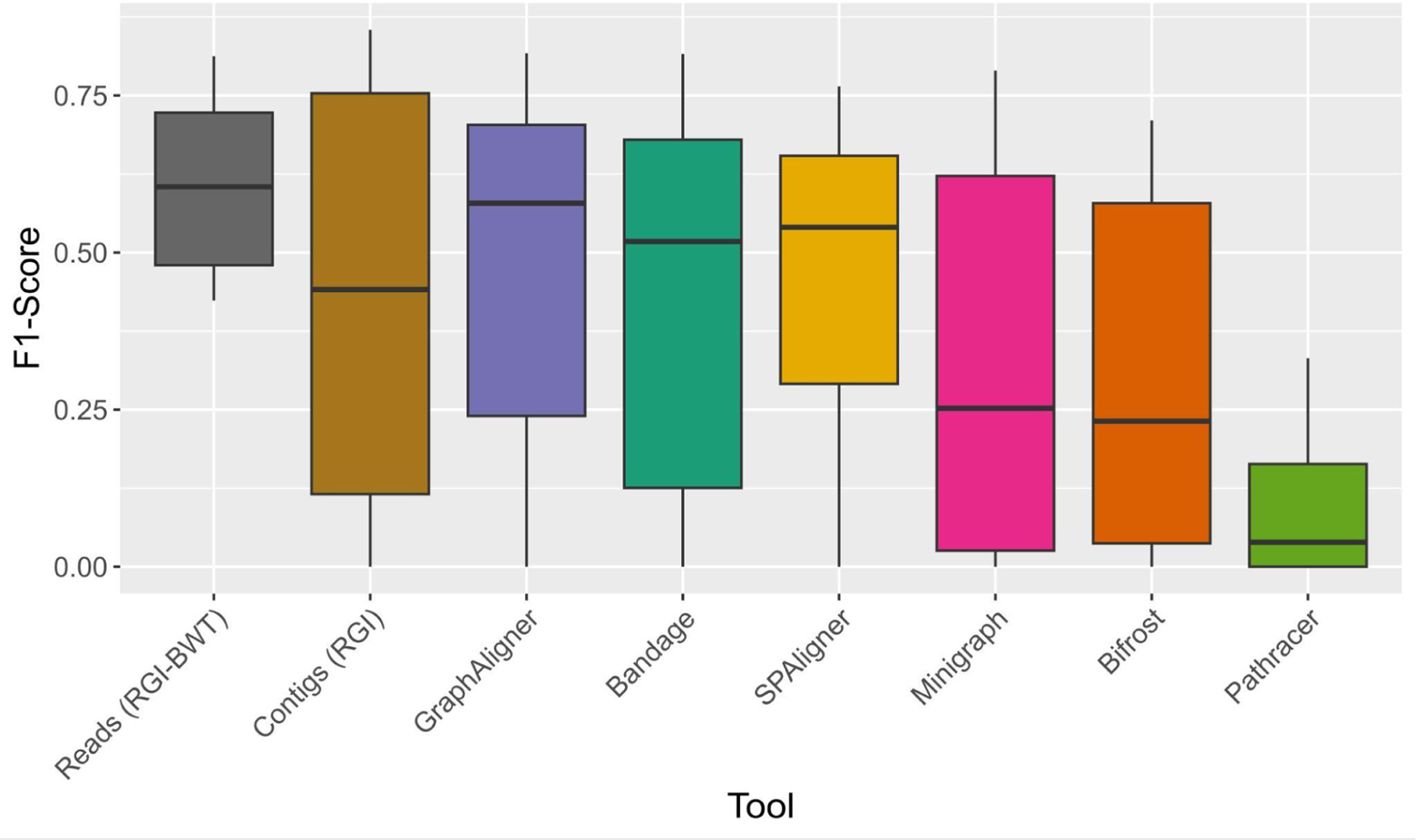
Boxplot of F_1_-scores of graph querying, contig, and read based tools in identifying the presence of CARD protein homolog model AMR genes from high complexity (6104 taxa), mobile element-enriched (30 taxa), low complexity (29 taxa) and ZymoBIOMICS (10 taxa) long and short-read datasets. Summary statistics shown are the median, interquartile range (IQR), the whiskers represent 1.5 • 25th and 75th percentiles.

AMR genes conferring distinct phenotypes can differ by just one nucleotide making the identification of specific alleles challenging which is why we also calculated the precision and recall of AMR gene families sub-clustered into groups that share at least 90% identity (6). Across all datasets and tools, mean precision increased from 0.69 to 0.75 when analysed at the family cluster level while recall remains similar (Figure 3).

**Figure 3.**
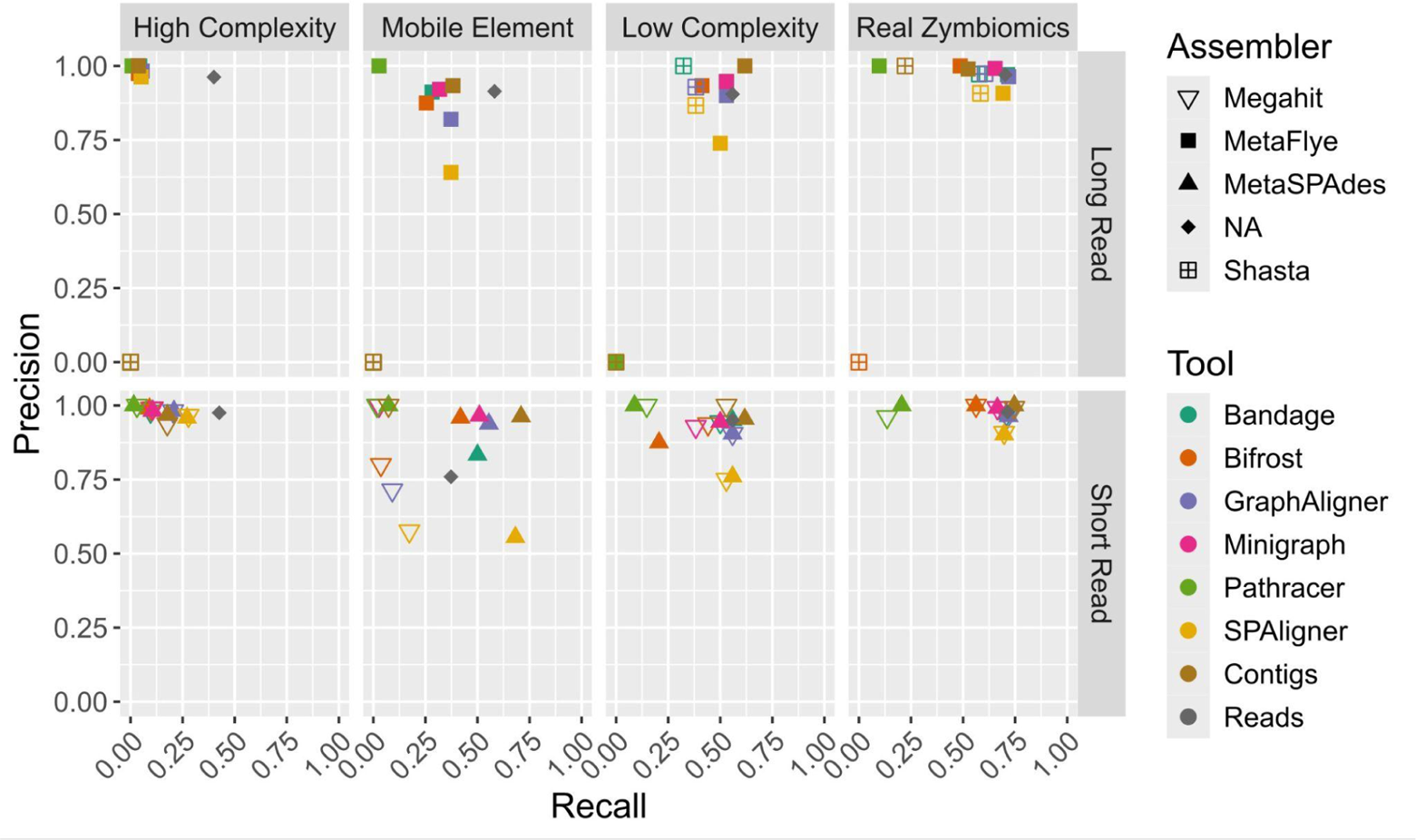
Precision vs. recall of graph querying, contig, and read based tools in identifying the presence of CARD protein homolog model AMR gene family clusters for high complexity (6104 taxa), mobile element-enriched (30 taxa), low complexity (29 taxa) and ZymoBIOMICS (10 taxa) long and short-read datasets assembled using MetaSPAdes, MetaFlye, Megahit, and Shasta.

### Graph-based methods moderately outperform contig AMR gene querying in complex datasets

To put the performance of graph-based methods in the context of more widely used methods, read-based AMR gene identification and assembled contig-based AMR gene identification were evaluated as well. Long- and short-read analyses on the high complexity graph identified more AMR genes than all other methods but were less precise (Figure 1, Figure 3). Long- and short-read based analyses on the low complexity and ZymoBIOMICS graphs were comparable in precision and recall to the contig-based analyses (Figure 1, Figure 3). Contig-based analyses performed well across all datasets with only select graph-based tools outperforming contigs in high complexity, mobile element-enriched long-read, and ZymoBIOMICS long-read graphs (Figure 1, Figure 3). For example, in the short-read high complexity MetaSPAdes graph, RGI ran on contigs outperformed all but two graph querying methods, correctly identifying the presence of 181 of 1348 AMR genes while GraphAligner identified 219 of 1348 and SPAligner identified 264 of 1348 AMR genes (Table S1).

### Fewer AMR genes are detected in high-complexity long-read metagenomes than short-read metagenomes across simulated read depths

Long-read simulated and mock metagenomes were also evaluated to assess the suitability of long-read metagenomics for AMR surveillance. Querying assembled contigs led to reduced recall of AMR genes in the high complexity, mobile element-enriched, and ZymoBIOMICS datasets as compared to read mapping (Figure 1). This is similar to the previously observed poor recall of AMR genes from short-read metagenomic contigs (7). Long-read mapping led to a similar recall distribution as short-read mapping with the exception of the mobile element-enriched dataset where long-read mapping led to 0.55 recall for the long-read dataset and 0.32 for the short-read dataset (Figure 1, Table S1). Across graph querying tools, long-read recall was similar to the equivalent short-read dataset at lower sequencing depths except for the high complexity dataset where short-reads led to higher overall recall of AMR genes (Figure 1).

### Performance is reduced as read depth decreases only for more complex metagenomes

Datasets were subsampled to investigate the effect of read depth on AMR gene presence identification. As read depth increases in the high complexity and mobile element-enriched datasets, F_1_-score increases as well, meaning that more AMR genes were able to be correctly identified in the graphs (Figure 4). Conversely, even the lowest read subsamples (25%) of the low complexity and ZymoBIOMICS datasets do not have a lower F_1_-score suggesting that all AMR genes present were sampled at a high relative depth in these metagenomes. Contig-based AMR gene identification suffered similar performance reductions as reads were subsampled (Figure 4).

**Figure 4.**
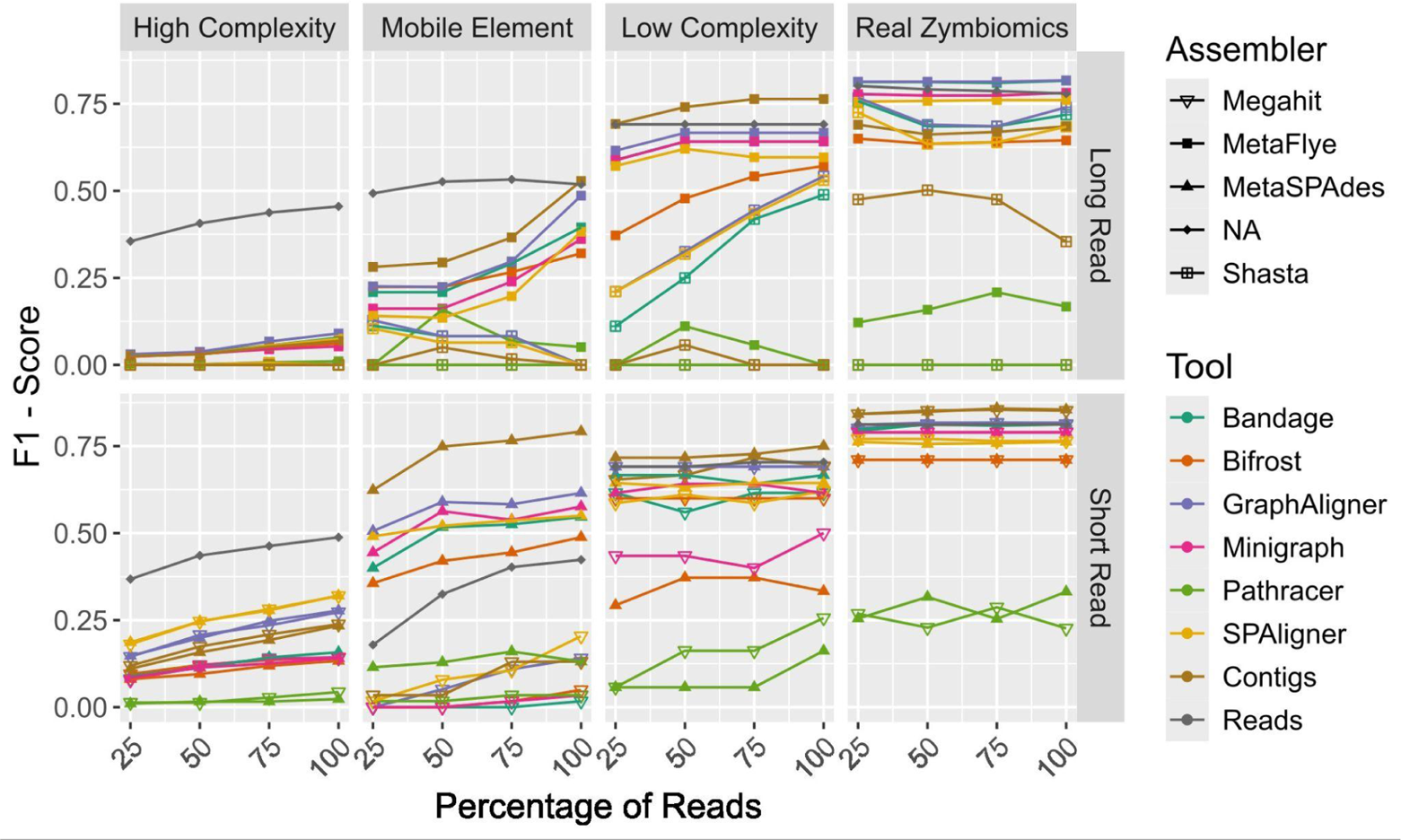
F_1_-score vs. read subsamples of graph querying, contig, and read based tools in identifying the presence of CARD protein homolog model AMR genes for high complexity (6104 taxa), mobile element-enriched (30 taxa), low complexity (29 taxa) and ZymoBIOMICS (10 taxa) long and short-read datasets assembled using MetaSPAdes, MetaFlye, Megahit, and Shasta.

### Graph- and contig-based AMR gene querying can be combined to maximise precision and recall

To assess the ability of graph-based and contig based methods to work synergistically to identify genes with genomic context, the results of RGI on the assembled contigs as well as GraphAligner on the assembly graph were combined. The combined analysis of assembly graphs and contigs resulted in higher recall in all data types except the low complexity long-read simulation (Figure 5). In the high complexity short-read and long-read datasets recall was increased from 0.13 to 0.21 and from 0.036 to 0.050 respectively using a combined graph and contig analysis relative to contigs alone (Figure 5). Similar recall improvements were found for mobile element-enriched short-read (0.68 to 0.73) and long-read (0.37 to 0.45) data; however, this was combined with lowered precision (0.94 to 0.51 and 0.91 to 0.63 respectively) (Figure 5).

**Figure 5.**
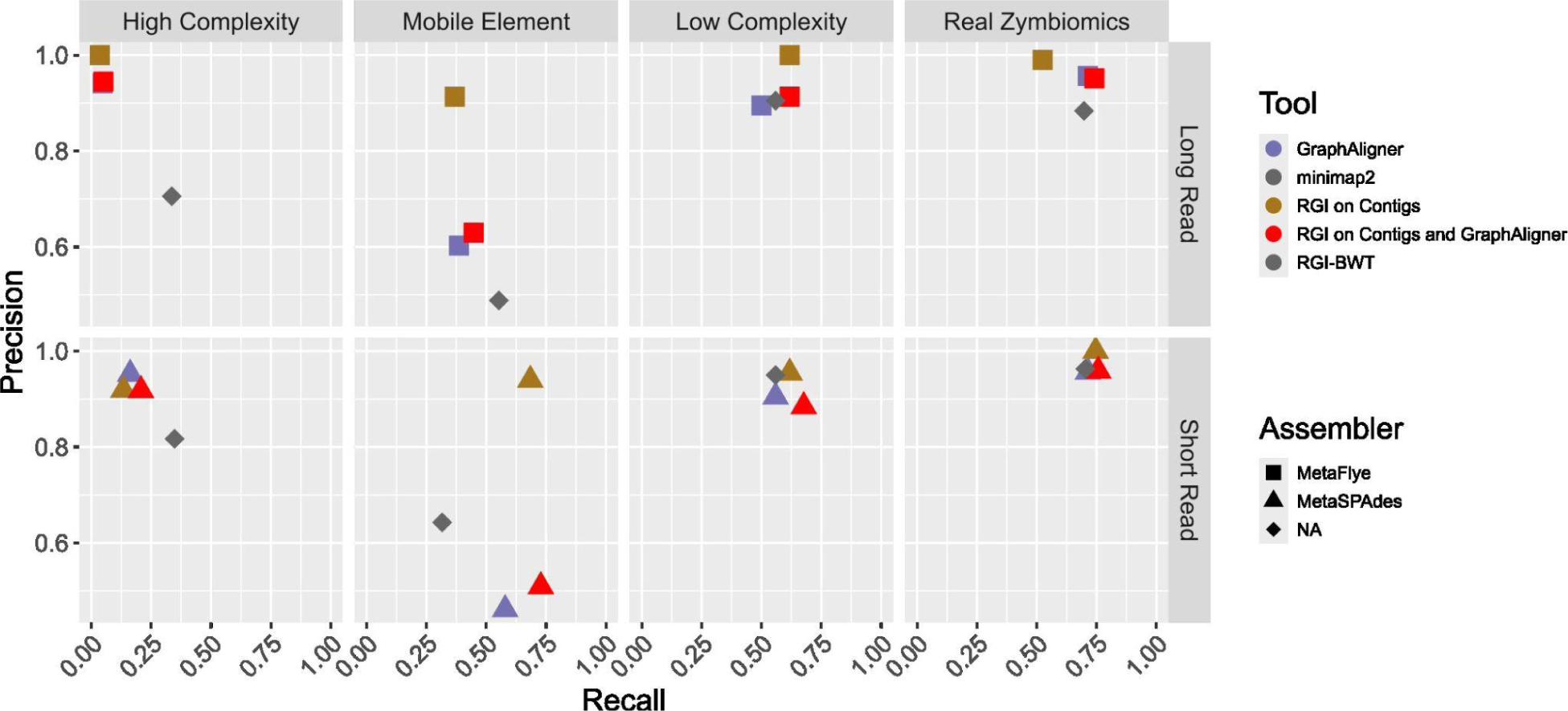
Precision vs. recall of GraphAligner, RGI run on assembled contigs, read based analysis, and combined RGI and GraphAligner results in identifying the presence of CARD protein homolog model AMR genes for high complexity (6104 taxa), mobile element-enriched (30 taxa), low complexity (29 taxa) and ZymoBIOMICS (10 taxa) long and short-read datasets assembled using MetaSPAdes, MetaFlye, Megahit, and Shasta.

## Discussion

### GraphAligner outperforms other tools for the detection of AMR genes in metagenomic assembly graphs

This study aimed to evaluate the suitability of BLAST (Bandage and GraphAligner), BWT (SPAligner via BWA-MEM2), K-mer (Bifrost and Minigraph), and HMM (Pathracer) based graph querying tools to the task of querying metagenome assembly graphs for the presence of AMR genes. In general, BLAST-based tools performed most strongly, specifically GraphAligner (Figure 2). Computationally, GraphAligner was not the fastest or the most memory efficient; however, it was only 3.22 minutes (193.4 s) slower and 1.3 GB more memory intensive on average than the top performing Bandage and Bifrost respectively. It should be noted that compared to the computational requirements of metagenomic assembly itself (required to infer the assembly graph) most querying methods were relatively efficient (Figure S1, Figure S2). GraphAligner had the highest F_1_-score, a harmonic mean of precision and recall, for all datasets except the short-read high complexity metagenome simulation (Figure 1, Figure 2). On this dataset, SPAligner identified the most AMR genes and should be considered by users interested in maximising their sensitivity in a high complexity metagenome. However, SPAligner was both more computationally demanding and produced output that was more challenging to parse for this task (Figure 1, Figure 3, Figure S1).

### Querying assembly graphs increases recall of AMR genes relative to contig-based analyses but only for high complexity metagenomes

Querying the metagenomic assembly graph has been proposed as an opportunity to identify AMR genes that fail to assemble into contigs while preserving the opportunity to investigate gene context (7, 9). On the high complexity metagenome, just GraphAligner and SPAligner were able to correctly identify more AMR genes than RGI run on the metagenomically assembled contigs (Figure 1, Figure 3). This pattern was not consistent across other complex samples (e.g., GraphAligner outperformed RGI on the mobile-element long-read metagenome MetaFlye graph but not on the metaSPAdes graph). On the taxonomically simpler low complexity (n=29) and ZymoBIOMICS (n=10) short-read datasets, the contig-based analysis outperformed all graph-based tools assessed (Figure 1, Figure 3). The read subsampling analysis suggests that this may be attributable to insufficient read-depths for higher complexity datasets (Figure 4). Together, these results suggest that graph-based tools can improve detection of AMR genes in complex samples but are unnecessary in simpler/well-sequenced samples. Given the relative complexity and mobile-element enrichment of priority surveillance environments, such as municipal wastewater, this suggests graph-based methods could represent an improvement on current surveillance methods (3).

### Long-read metagenomics faces similar AMR gene identification challenges as short-read metagenomics

Analysis of contigs assembled from long-read metagenomes identified fewer AMR genes than read- or graph-based analyses (Figure 1). This demonstrates that, similar to short-read assembly, the process of resolving contigs from long-read assembly graphs leads to fewer identifiable AMR genes (7). Precision and recall were better for datasets assembled with MetaFlye rather than Shasta. Unlike short-reads, assembler choice may have a larger impact when identifying AMR genes from long-read metagenomes (Figure 1). Long-read metagenomics performed worse, across all AMR gene identification methods tested, than short-read metagenomics for the high complexity dataset (Figure 1). Read subsampling analyses suggested that, similarly but more severely than short-reads, this may be a consequence of insufficient long read-depths for complex samples (Figure 4). However, even well-sampled low complexity metagenomics (e.g., ZymoBIOMICS dataset) did not result in higher AMR gene recall for long-reads than short-reads (Figure 1, Figure 4). This suggests that there are still unsolved technical challenges in accurately identifying AMR genes from long-read data.

### Graph- and contig-based AMR gene detection can be combined to maximise recall

The metagenomic assembly tools used in this study output both the assembled contigs as well as the assembly graph. If it is the goal of a metagenome analysis to identify as many real AMR genes as possible, analysing the assembly graph may lead to 55% to 75% increases in the number of genes correctly identified relative to contig-analyses alone (Figure 5).

In high complexity metagenomes it is unlikely that graph- or contig-based methods (or both combined) will ever be as sensitive as read-based methods. Metagenomic enrichment approaches such as bait-capture and selective sequencing can also improve AMR gene recovery but contribute additional biases and are unlikely to enrich the potentially unknown genes surrounding a target AMR gene (19–21). Recovery of this wider genomic context is a fundamental challenge with the fragmented DNA and insufficient sampling depth common to complex metagenomic samples. Despite the overall low gene recall of AMR genes with graph- or contig-based analyses (even when combined), for those genes that are detected the genomic context information greatly increases their utility to public health surveillance and risk-assessment (8–10).

### Study limitations

This study evaluated the ability of graph alignment tools to accurately identify the presence or absence of an AMR gene in a metagenomic assembly graph, not the quantity of each AMR gene present in the dataset. When the precision and recall of the quantity of AMR genes identified was calculated, recall dropped substantially, especially for the complex datasets, suggesting that the collapse of redundant sequences in the assembly graph impedes simple quantification (Figure S3, Figure S4). Future work should explore the performance of k-mer coverage-based metrics for the accurate estimation of gene copy number from graphs. It should also be noted that we did not confirm the assembly was error free for each AMR gene. Assembly errors will occur at varying rates across the datasets and this risk should be taken into account when analysing AMR gene context (22). Finally, this study used the final, error corrected, and condensed assembly graph output from each assembler which is the final step before paths are walked to generate contigs (23–26). Intermediate assembly graph analysis could potentially allow for further increases in AMR gene recall.

## Conclusion

Identifying as many AMR genes, and their surrounding genomic contexts, from metagenomic sequencing data is of critical value to public health AMR surveillance. This study demonstrates that for taxonomically complex microbiomes, BLAST-based assembly graph analyses can lead to increases in AMR gene recall especially when combined with contig-based analyses. Additionally, this study demonstrates that despite some key advantages, current long-read metagenomic sequencing may face similar challenges as short-read metagenomics in relation to AMR surveillance. These results will inform the integration of further graph-based AMR gene identification into metagenomic surveillance pipelines to better mitigate the burden of AMR on human and animal health.

## Methods

### Dataset and metagenome simulation

Three simulated and one mock microbial community were used for this study, a taxonomically high complexity sample, a mobile element-enriched sample, a taxonomically low complexity sample, and finally the simulation independent ZymoBIOMICS mock community standard (12). The high complexity metagenome was simulated based on the taxonomic composition of a real agricultural soil metagenome (NCBI SRA Accession SRR5456987) from the Earth Microbiome Project (27). The taxonomic composition and relative abundance were obtained using Kraken2 in unique mode (v2.1.3) and Bracken (v2.9) respectively (28–30) with the NCBI RefSeq Standard database (6/5/2024) (31). This resulted in the identification of 6104 unique taxa in this microbial community (Table S2). To simulate the low complexity metagenome the composition and relative abundances were inferred from a pediatric healthy lung sample resulting in 29 unique taxa (32). Finally, the mobile element-enriched metagenome was based upon a simulation used in a previous investigation of metagenomic assembly which consists of 30 taxa rich in plasmids and genomic islands (7). For each of these simulations, the corresponding genome assemblies were downloaded from NCBI using the NCBI datasets tool (v15.24.0). Nanosim (v3.1.0) was then used to simulate 3 million read ONT long-read metagenomes using the ZymoBIOMICS even community pre-trained model and allowing for chimeric contigs for all samples (12, 33). Long-read sequencing depth was based on previous studies evaluating the suitability of long-reads for metagenomics as well as the ZymoBIOMICS mock microbial community standard which ranged the sequencing of metagenomes from 1 to 5 million reads (12, 34). Read depth is not changed as sample complexity increases as extracting sufficient DNA is often challenging from complex sample matrices (35). InSilicoSeq (v1.6.0) was used to simulate 60 million Illumina MiSeq reads for the high complexity metagenome and 30 million Illumina MiSeq reads for the low complexity and mobile element-enriched metagenomes (36). Short-read depths are the same as the quantity of prokaryotic short-reads from the original soil, lung, and mobile element-enriched metagenomes these simulations are based upon (7, 27, 32). The simulation independent dataset is the ZymoBIOMICS mock community standard consisting of genome assemblies of each species in the mock community, Illumina MiSeq, and ONT GridIon sequencing of the mock microbial community (12).

### Baseline Antimicrobial Resistance Gene Profile

To establish a ground truth, the RefSeq assemblies used in the metagenome simulations as well as the genome assemblies of the ZymoBIOMICS mock microbial community were annotated for protein homolog model AMR genes using the Comprehensive Antibiotic Resistance Database (CARD v3.2.8) and Resistance Gene Identifier (RGI v5.2.0) excluding loose hits (12, 31, 37).

### Graph Construction

Simulated and mock metagenome reads were quality filtered then assembled. For short-reads, fastp (v0.23.4) was used with default settings followed by assembly with metaSPAdes (v3.15.5) and megahit (v1.2.9) (23, 24, 38). For long-reads, filtlong (v0.2.1) was used to filter out reads shorter than 1000bp and the worst 10% of reads followed by assembly with Flye (v2.9.2) in metagenome and pre-corrected reads mode as well as Shasta (v0.12.0) (26, 39, 40). Of the long-read assemblers tested, Flye and Shasta produced graphs compatible with the largest number of graph querying tools being analysed (Table S3). Megahit fastg assembly graphs were produced for the largest k-mer iteration (k = 141) and converted to GFA format using Bandage (v0.8.1).

### Graph Querying Tools Assessed

All assembly graph-query capable tools were assessed for their accuracy, sensitivity, recall, and computational resource demand in identifying and locating CARD protein homolog model genes (v3.2.8) within GFA-formatted assembly graphs. The Bandage (v0.8.1) querypaths function and GraphAligner (v1.0.17b) were used with default settings (16, 18). SPAligner, a submodule of SPAdes (v3.15.5), was used with the default configuration parameters as well as the CARD protein homolog model nucleotide sequences converted to fastq format with perfect quality scores using Biopython (v1.83) as SPAligner does not support fasta format (13, 41). The Bifrost (v1.3.0) -P setting was used to output the ratio of found k-mers for each CARD protein homolog model query. Bifrost, as natively implemented, does not locate any of the k-mers aligned to the graph. However, in the Bifrost UnitigMap data structure, the location of each k-mer is stored. To access the location of found k-mers, Search.ttc was modified to output the first k-mer of the unitig containing a pseudoalignment as well as the first k-mer of the pseudoalignment (14). Minigraph (v0.20) was used with CARD protein homolog model nucleotide sequences in fasta format; however, Minigraph does not support sequence graphs with segment overlaps as is the case with SPAdes, Megahit, and Shasta assembly graphs. To remove the overlaps in the SPAdes and Megahit assembly graphs, the remove_all_overlaps() function from Unicycler (v0.5.0) was used and modified to prevent the trimming of segments to zero (15, 42). Shasta graph segments contain variable length overlaps and were not able to be trimmed and therefore excluded from Minigraph analyses. Shasta overlap graphs were also incompatible with Bifrost and Pathracer (26). This graph processing was included in the total computational resource demand for Minigraph. Pathracer (v3.16.0) requires the construction of Hidden Markov Models (HMMs) to be used as input. To do this CARD protein homolog model amino acid sequences were first grouped by AMR gene family then clustered into sub-families of 90% amino acid identity using MMseqs2 (v15.6) (43). These 90% identical sub-families were then aligned with MUSCLE (v5.1) and used to build HMMs using HMMER (v3.4) (44, 45). Pathracer is implemented for AMR gene detection in a pipeline called graphamr which further processes Pathracer results to deal with assembly graph overlaps and other sequence graph properties (46). To assess Pathracer we implement the same approach as the graphamr pipeline following Pathracer graph querying which includes open reading frame (ORF) identification in six frames and ORF clustering at 90% identity using MMseqs2 (v15.6), followed by annotation of those sequences using RGI (v5.2.0) (37, 43, 46). To ensure a fair comparison only the Pathracer graph querying itself is included in the computational resource demand measurement. Computational resource usage was recorded using Nextflow’s trace function (47).

### Read- and contig-based AMR gene querying

To put graph-querying methods in the context of existing methods to query metagenomic data for AMR genes, simulated and ZymoBIOMICS reads as well as their assembled contigs were queried for the presence of CARD protein homolog model genes (v3.2.8). Short-reads were queried using RGI-BWT (v5.2.0) which used KMA to map reads onto the redundant CARD database (6, 37). For the long-reads, minimap2 (v2.28) was used to align the reads to the CARD database followed by samtools (v1.2.0) to calculate coverage (48, 49).

### Data Analyses

Graph querying tools were run on a computer with an Intel(R) Xeon(R) Silver 4114 CPU with 40 cores and 502 GB of memory with an Ubuntu (v22.04) operating system. Data analyses and visualisations were done in R (v4.3.1).

Minigraph, Bifrost, Bandage, Graphaligner, and SPAligner do not assign one hit per locus (as is the case with Pathracer/graphamr implementing an ORF calling step) but do report the coordinates of each hit. A script (subsampled_hit_region.R) using the coordinates of each hit was written to identify each distinct hit locus. This includes hits that are only on one segment of the assembly graph as well as hits that span multiple segments (Figure S5). As CARD contains many near identical sequences (i.e., different alleles of the same AMR gene) there were many hits on a single hit locus. To determine which hit was best, the following criteria were applied in order: the highest query coverage, highest percent identity, and longest hits were chosen to be the best hit for a given hit locus. In cases where multiple hits satisfied the above conditions, the hit with highest lexical order CARD ARO accession was chosen. The SPAligner output failed to delimit the alignment length of some multi-segment alignments which required post-processing verification using the graph segment lengths.

Precision and recall were calculated to evaluate tool performance as compared to the baseline AMR gene profile. Precision and recall of binary AMR gene presence/absence was calculated in addition to the precision and recall of total AMR gene quantity. Binary gene presence/absence is likely more reflective of the true precision and recall of the graph query tools as assembly graphs collapse repeats into single high coverage segments making coverage a more appropriate indicator of metagenomic AMR gene copy number (50).

### Data summary

All analyses described above can be reproduced and investigated further using scripts found in the following GitHub repository (https://github.com/maguire-lab/metagenomic_graph_querying_benchmarking), Zenodo code repository (https://doi.org/10.5281/zenodo.16987655) and OSF data repository containing the assembly graphs and contigs DOI 10.17605/OSF.IO/QJX54 (https://osf.io/qjx54/)

## Supplemental Figure and Table Captions

Table S1. Quantities of correctly identified AMR genes and quantities of identified AMR genes per dataset.

Table S2. Taxa and relative abundances identified using Kraken2 in unique mode (v2.1.3) and Bracken (v2.9) from a soil metagenome from the Earth Microbiome Project (NCBI SRA Accession SRR5456987).

Table S3. Summary of long-read assemblers and their suitability to produce graphs that can be readily queried for the presence of AMR genes.

**Figure S1.**
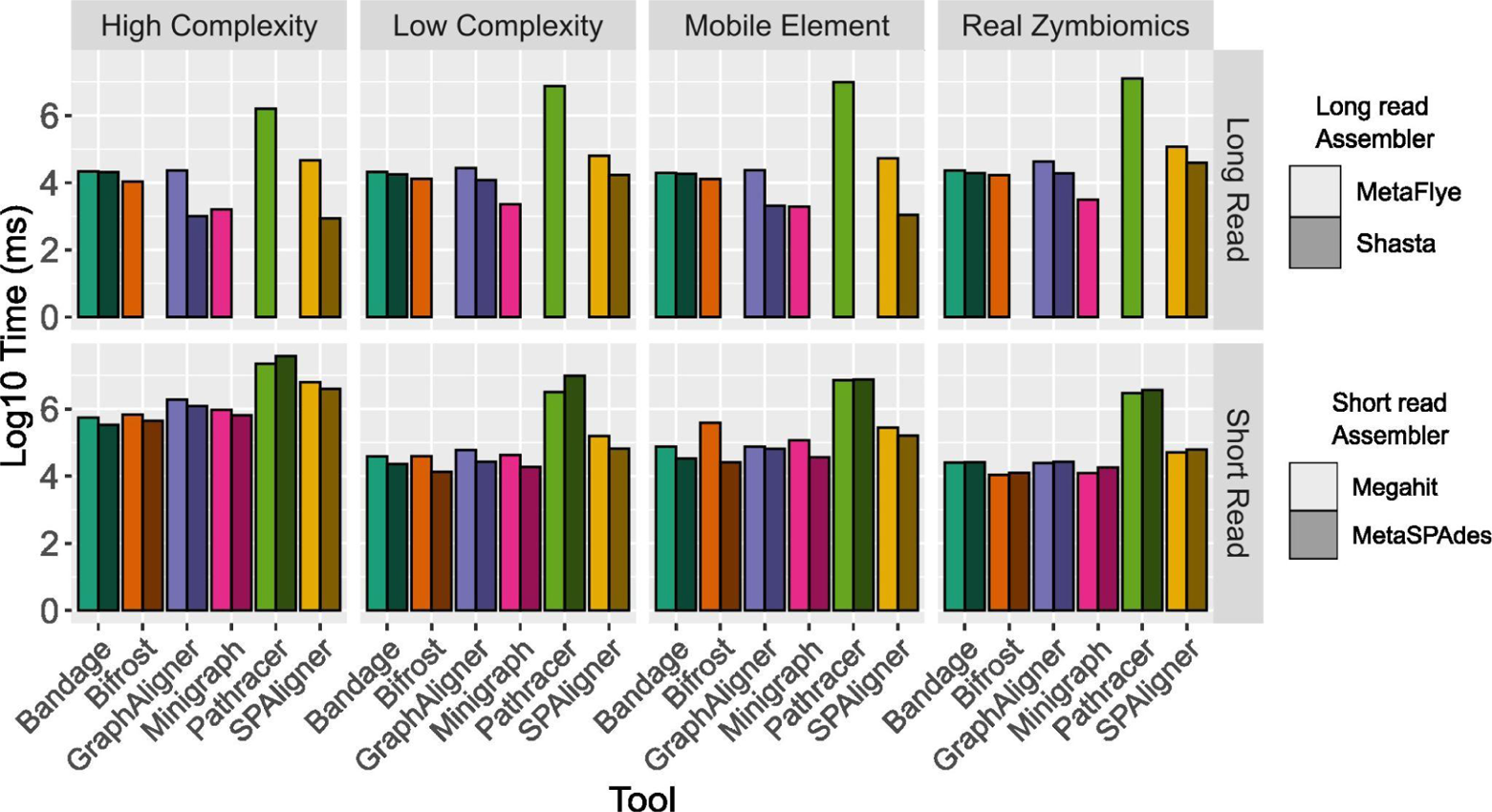
Runtimes (log ms) for graph querying tools to query for AMR genes in the CARD protein homolog model database on high complexity (6104 taxa), mobile element-enriched (30 taxa), low complexity (29 taxa) and ZymoBIOMICS (10 taxa) long and short-read datasets assembled using MetaSPAdes, MetaFlye, Megahit, and Shasta.

**Figure S2.**
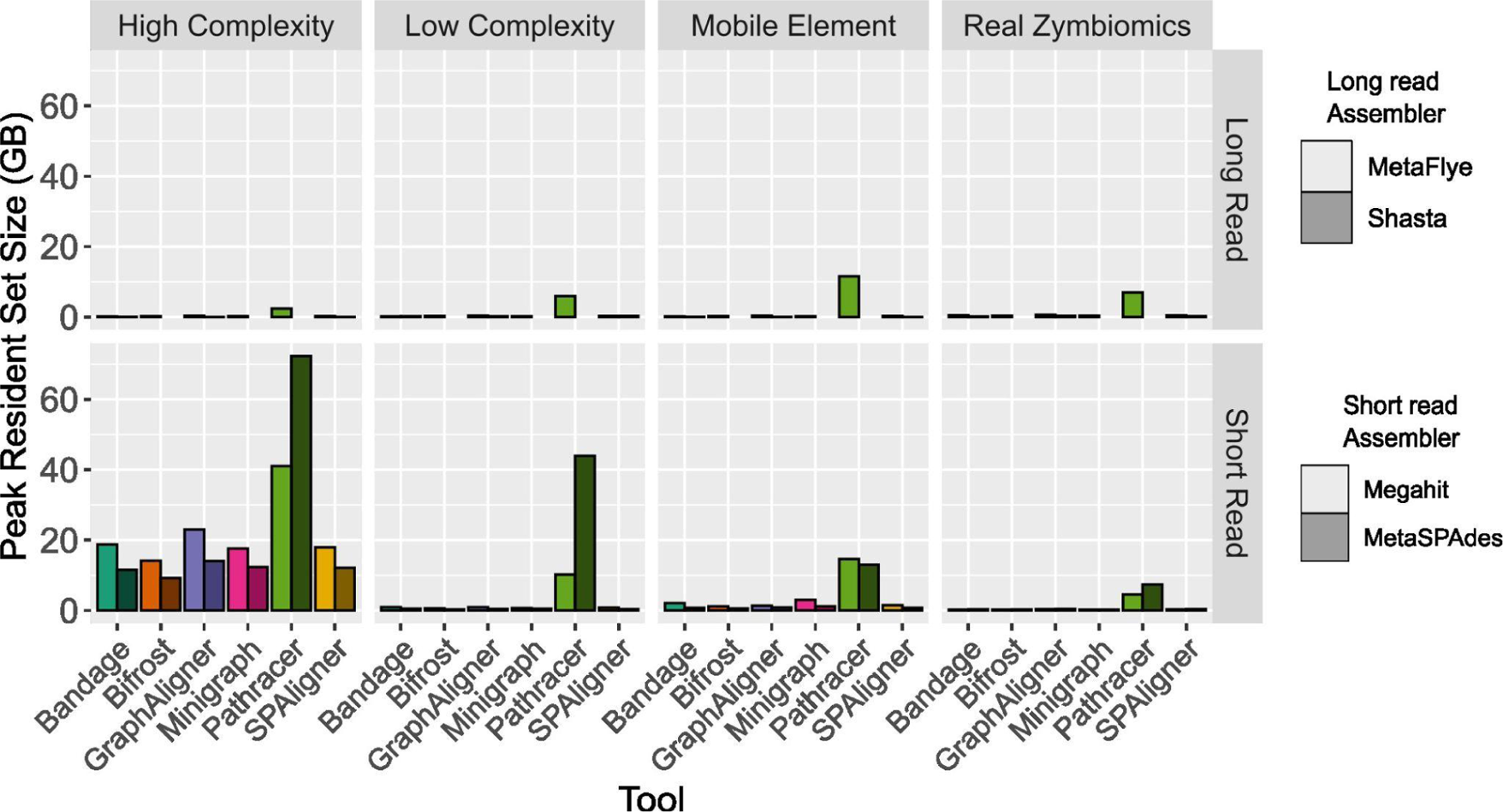
Peak memory usage (resident set size) for graph querying tools to query for AMR genes in the CARD protein homolog model database on high complexity (6104 taxa), mobile element-enriched (30 taxa), low complexity (29 taxa) and ZymoBIOMICS (10 taxa) long and short-read datasets assembled using MetaSPAdes, MetaFlye, Megahit, and Shasta. Peak memory usage when assembling the reads was 17 GB for ZymoBIOMICS, 151 GB for low complexity, 446 GB for mobile element enriched, and 90 GB for high complexity datasets.

**Figure S3.**
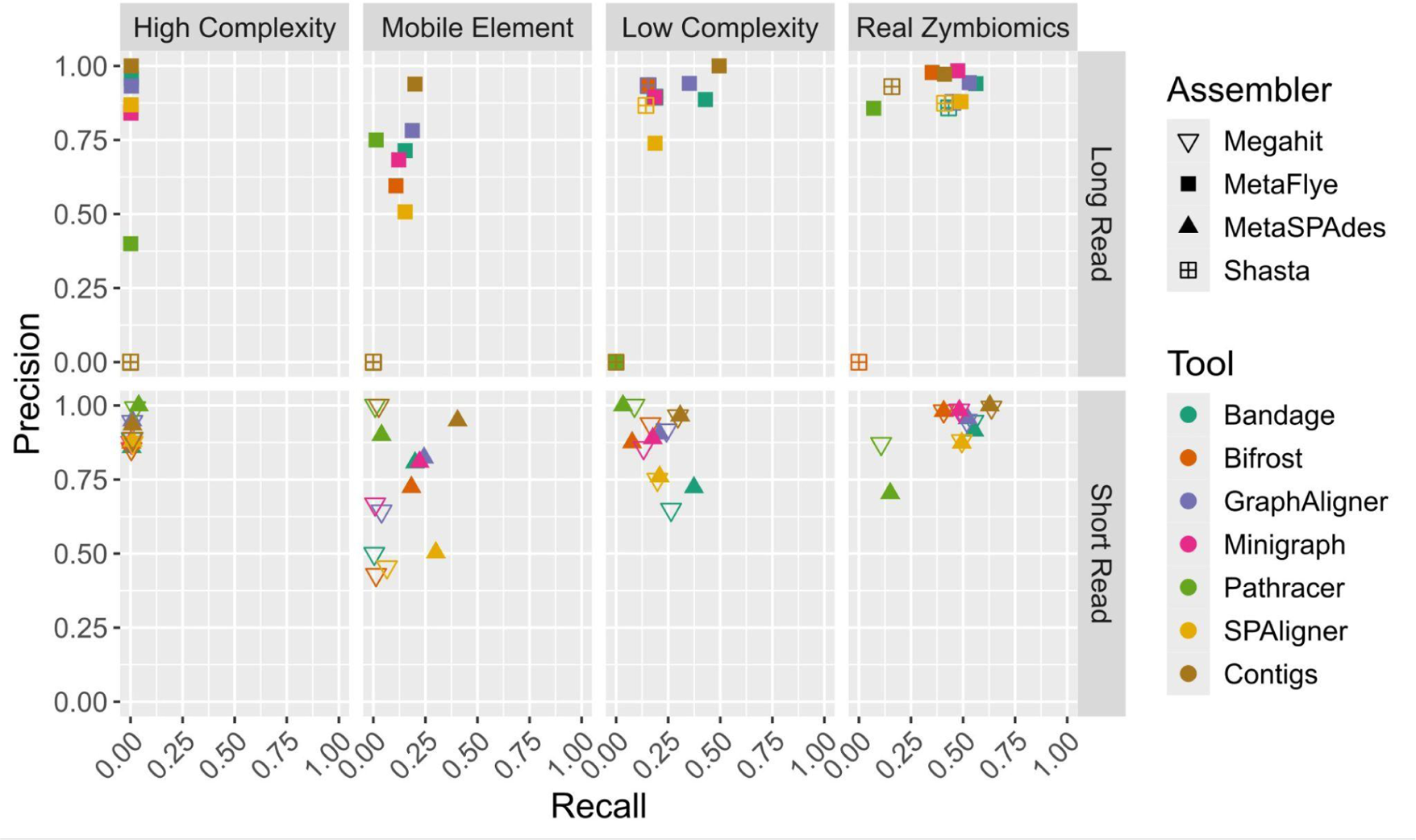
Precision vs. recall of graph querying, contig, and read based tools in quantifying CARD protein homolog model AMR genes for high complexity (6104 taxa), mobile element-enriched (30 taxa), low complexity (29 taxa) and ZymoBIOMICS (10 taxa) long and short-read datasets assembled using MetaSPAdes, MetaFlye, Megahit, and Shasta.

**Figure S4.**
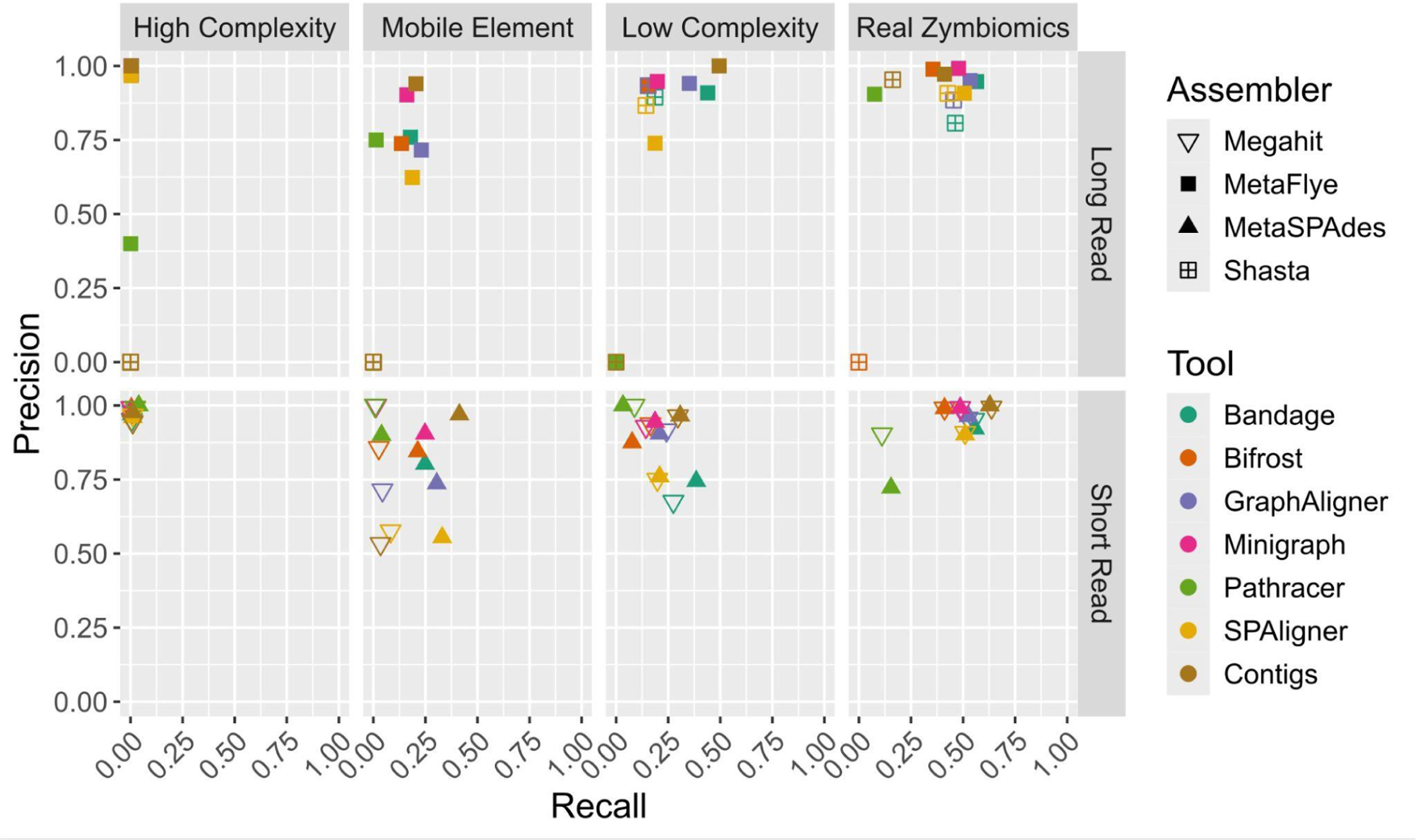
Precision vs. recall of graph querying, contig, and read based tools in quantifying CARD protein homolog model AMR gene family clusters for high complexity (6104 taxa), mobile element-enriched (30 taxa), low complexity (29 taxa) and ZymoBIOMICS (10 taxa) long and short-read datasets assembled using MetaSPAdes, MetaFlye, Megahit, and Shasta.

**Figure S5.**
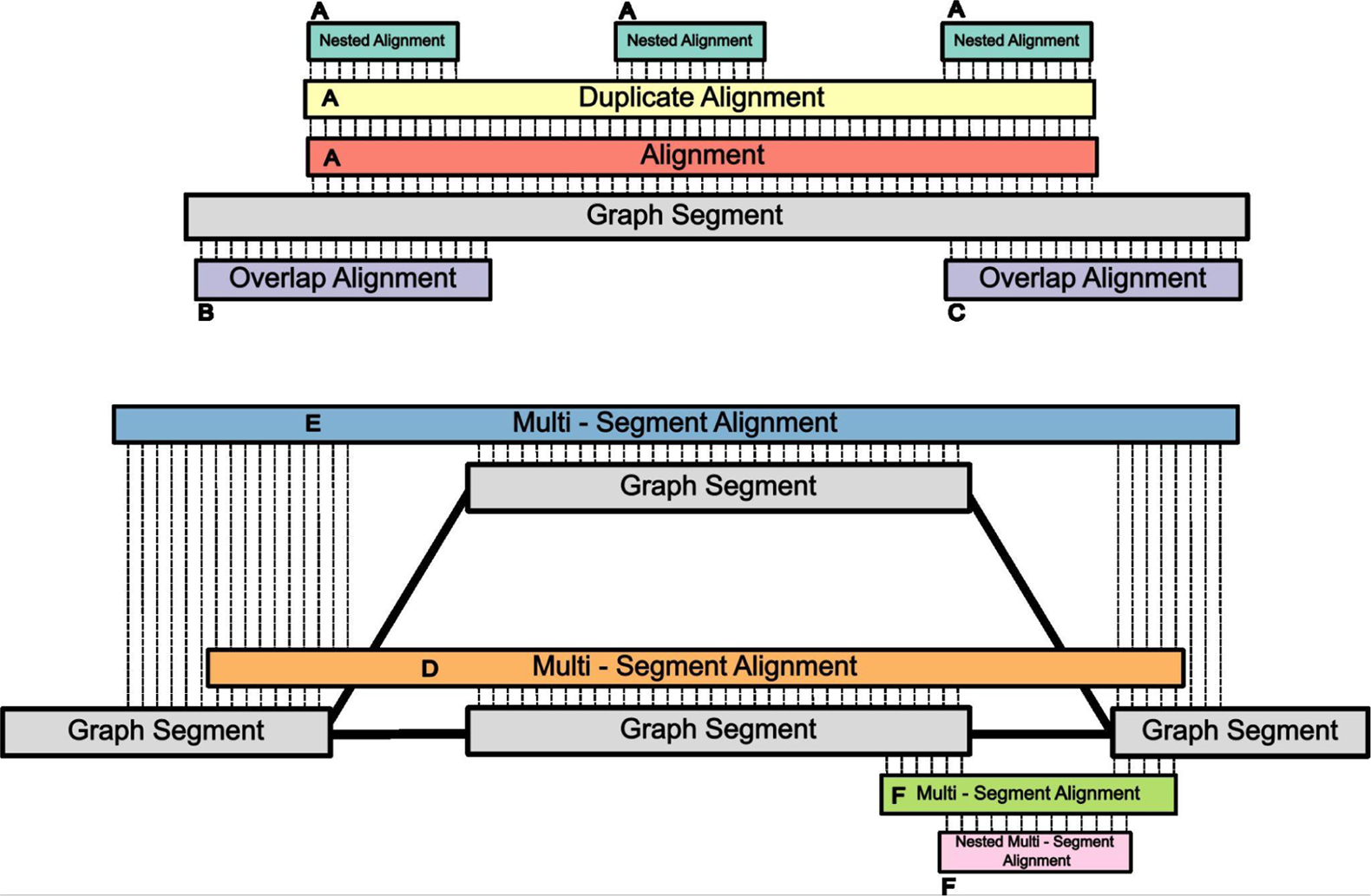
Graph alignment scenarios and which types of graph alignments get assigned within the same hit locus or in distinct hit loci. Hit locus A covers single segment hits, duplicate hits in the exact locus, and smaller nested hits within them. Overlapping hits are assigned distinct loci (B and C). Multi segment hits are assigned a distinct locus if they align to different segments (D, E, and F). However, as seen in locus F, when there is a nested multi-segment alignment, they are assigned to the same hit region. Once hit loci are identified, the best hit for each locus is selected.

